# Automated deep lineage tree analysis using a Bayesian single cell tracking approach

**DOI:** 10.1101/2020.09.10.276980

**Authors:** Kristina Ulicna, Giulia Vallardi, Guillaume Charras, Alan R. Lowe

## Abstract

Single-cell methods are beginning to reveal the intrinsic heterogeneity in cell populations, which arises from the interplay or deterministic and stochastic processes. For example, the molecular mechanisms of cell cycle control are well characterised, yet the observed distribution of cell cycle durations in a population of cells is heterogenous. This variability may be governed either by stochastic processes, inherited in a deterministic fashion, or some combination of both. Previous studies have shown poor correlations within lineages when observing direct ancestral relationships but remain correlated with immediate relatives. However, assessing longer-range dependencies amid noisy data requires significantly more observations, and demands the development of automated procedures for lineage tree reconstruction. Here, we developed an open-source Python library, *btrack*, to facilitate retrieval of deep lineage information from live-cell imaging data. We acquired 3,500 hours of time-lapse microscopy data of epithelial cells in culture and used our software to extract 22,519 fully annotated single-cell trajectories. Benchmarking tests, including lineage tree reconstruction assessments, demonstrate that our approach yields high-fidelity results and achieves state-of-the-art performance without the requirement for manual curation of the tracker output data. To demonstrate the robustness of our supervision-free cell tracking pipeline, we retrieve cell cycle durations and their extended inter- and intra-generational family relationships, for up to eight generations, and up to fourth cousin relationships. The extracted lineage tree dataset represents approximately two orders of magnitude more data, and longer-range dependencies, than in previous studies of cell cycle heritability. Our results extend the range of observed correlations and suggest that strong heritable cell cycling is present. We envisage that our approach could be extended with additional live-cell reporters to provide a detailed quantitative characterisation of biochemical and mechanical origins to cycling heterogeneity in cell populations.

## INTRODUCTION

Individual cells grown in identical conditions within populations of either clonal or closely related origin often exhibit a highly heterogeneous proliferative behaviour^1^. Deciphering why and how cell heterogeneity is established, maintained and propagated over generations remains a key challenge. This is increasingly important in studying dynamic developmental processes involving the emergence of diverse cell types committed to different cell fates^2^, as well as in pathological scenarios.

Heterogeneity in cell populations may originate from deterministic or stochastic processes. For example, the observed distribution of cell cycle durations in a population of cells is governed either by stochastic processes, inherited in a deterministic fashion, or some combination of both^3–6^ (**Figure 1A**). Previous studies have shown poor correlations within lineages when observing direct ancestral relationships, but remain correlated with immediate relatives^4,7,8^. However, these studies have been performed using manually annotated data, which is laborious to acquire and limits the depth and statistical power to study more distant relationships amid noisy data.

**Figure 1.**
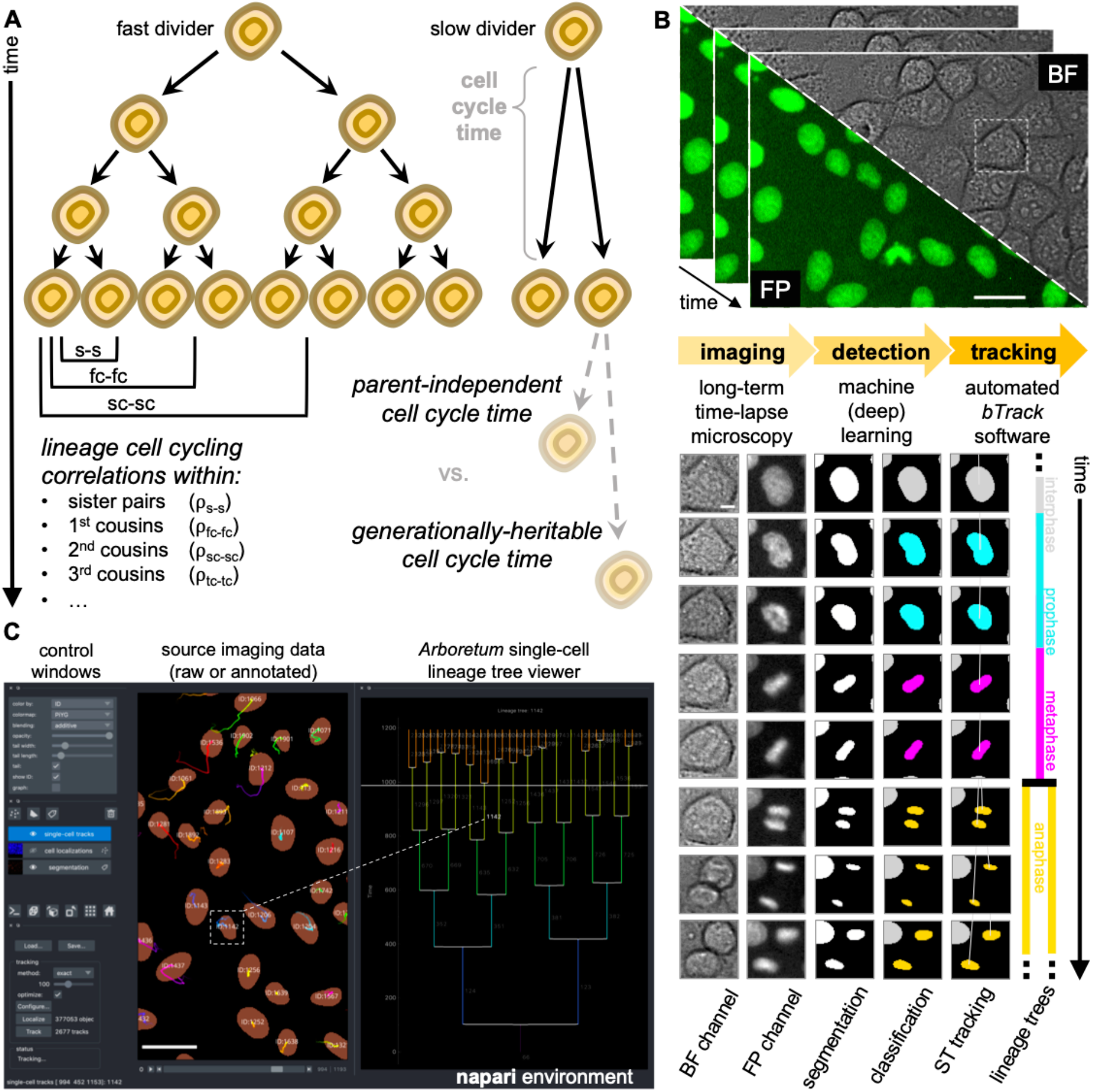
Overview of experimental and computational design. **(A)** We analyse heritability of cell cycle duration across multiple cell generations using automatically reconstructed multi-generational lineage trees. (**B)** Overview of the experimental *(data acquisition)* and computational *(data analysis)* workflow to address the stated biological problem. *(Top)* Sequential fields of view obtained by live-cell imaging experiments. Scale bar = 20μm. *(Bottom, left to right)* Fully automated, deep learning-based movie analysis consists of cell detection step, using information from bright-field and fluorescence time-lapse microscopy. Cells are localised using the segmentation and labelled according to their mitotic state by classification. Our software then reconstructs individual cell trajectories using this information and assembles the parent-children relationships into lineage tree representations. Scale bar = 5μm. BF, bright field; FP, fluorescent protein; ST, spatio-temporal. (**C)** A graphical user interface plugin is provided to enable users to interact with the imaging data and automated single-cell tracking and lineage analysis.

To increase the throughput of studies focussing on cell relationships within their reconstructed lineages, researchers need access to automated processing tools of large data quantities with single cell resolution^9^. The popularity of automated time-lapse microscopy has led to rapid growth in the volume and complexity of the data being acquired^10^. Until recently, one of the major challenges has been the segmentation of individual cells from live-cell images in ranging cell densities and fluorescence intensities. However, significant progress has been achieved in computer vision and machine learning approaches which enable robust segmentation and localisation of cells in these datasets^11–16^.

Unlike the cell detection step, connecting the single-cell observations from segmented image sequences over time into a biologically relevant trajectories, remains a challenging problem^1^. In addition to reconstructing trajectories over the lifetime of a single cell, it is also essential to correctly identify cell divisions and the relationships between cells to precisely reconstruct single-cell lineages. This is made more challenging in crowded conditions with high cell densities and diverse migration patterns.

Accurate tracking of individual cells, and reconstruction of cell lineages from raw microscopy images remains a major bottleneck and rate-limiting step in subsequent data analysis despite major efforts in this area^17–29^. To extract multigenerational lineages, it is often necessary to manually annotate ancestor/descendant relationships in graphical lineage tree representations. Monitoring cells captured over a period of a few days can easily demand for several weeks of experienced annotator’s dedicated time^19^ to manually reconstruct the trees which capture cell observations for about 10 generations. Even the most advanced cell tracking frameworks require some degree of user’s supervision (so-called semi-automated pipelines, extensively reviewed in^1,10,30^). Recent developments introduced automated approaches to detect and correct the tracking discrepancies^31^. However, the remaining high incidence of errors in the tracking data often requires additional human oversight to manually curate, or correct, the tracker outputs. This laborious, time-consuming and often error-prone task currently results in trade-offs being made between the minimum experimental replicates sufficient for reliable low-throughput analysis and maximum volumes of imaging data that researchers are capable of semi-manual processing.

To address this challenge, we developed *btrack* (Bayesian Tracker), an open-source, fast and robust Python library for unsupervised single-cell tracking of cell populations imaged using time-lapse microscopy (**Figure 1B-C**). We benchmark the performance of our library by computing multiple metrics for cell detection, tracking performance and fidelity of lineage reconstruction. Using our *btrack* software on a large dataset of long-term time-lapse imaging data, we demonstrate that we can recover cell cycle durations from single cell tracks, generate lineage trees, and determine intra- and inter-generational correlations in cell cycle duration. Our automated approach has enabled us to analyse 5,325 individual cell lineages arising from analysis of 20,074 single cells to yield two orders of magnitude more single-cell data than in previous studies of cell cycle heritability.

## RESULTS

To assess the heterogeneity and the heritability of cell-cycle duration in a population of cells, we sought to automatically reconstruct family trees from individual cells in long-term time-lapse movies. We first acquired a large dataset of 44 long duration time-lapse movies of MDCK wild-type cells in culture, which comprises 52,896 individual 2 channel images (brightfield and the nuclear marker H2B-GFP), containing approximately 250,000 unique cells. This dataset spans densities from single cells to highly confluent monolayers, providing a unique dataset to thoroughly test the computational framework.

To automate data analysis, we developed a fast, open source and easy-to-use cell tracking library to enable calculation of intermitotic durations and the capture of multi-generational lineage relationships. Our pipeline consists of three steps: (i) cell segmentation, (ii) cell state labelling, where progression of cells towards division is classified, and (iii) cell tracking with lineage tree reconstruction. The reconstructed trees contain information such as generational depth, parent and root (i.e. founder) cell identity, as well as labelling of leaf cells (those with no known progeny).

### Bayesian cell tracking approach

The first step is to robustly localize cells in the raw image data. Recent advances in deep learning have demonstrated the utility of convolutional neural networks in cell instance segmentation from microscopy images. Therefore, we created a U-Net^32^ to detect individual cells based on their histone fluorescence (**Figure 2A**), using a per-pixel weighted loss function to separate proximal cells. To train and validate the network, we manually annotated a dataset of 150 fluorescent images (130 training, 20 validation, comprising 56,362 cells) and trained the U-Net to output a binary segmentation mask, representing individual cell nuclei (**Supplementary information**). Subsequent localization of the centroid coordinates for every detected object in the binary classification mask provides the minimal input to our tracking algorithm. Whilst we utilized our own U-Net in the present study, we note that several other recently developed cell segmentation algorithms, such as DeepCell^14,15^, StarDist^12^ and Cellpose^13^, can also provide input data to the tracking pipeline, which is agnostic toward the segmentation method used for cell detection.

**Figure 2.**
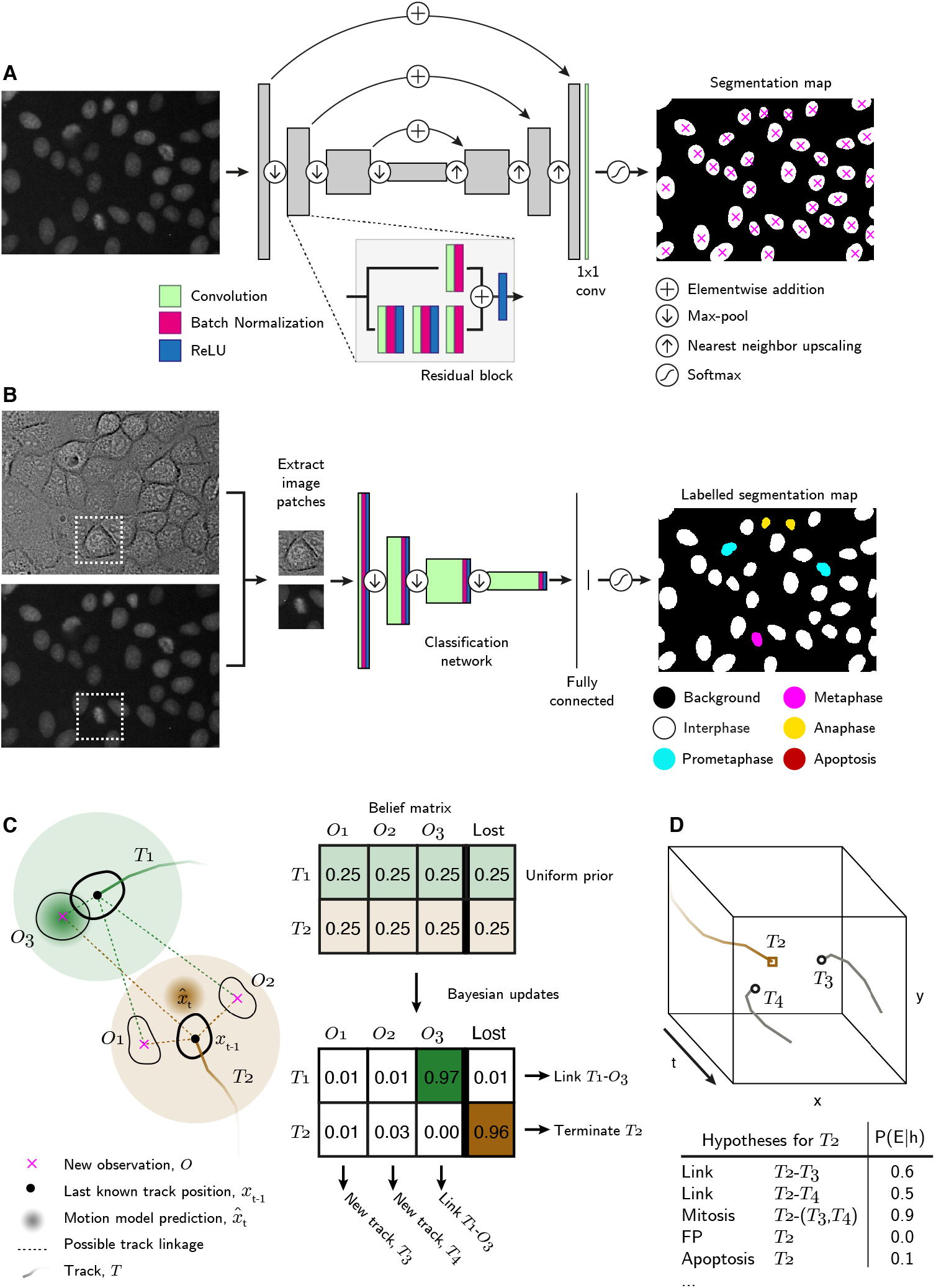
Single-cell tracking workflow. **(A)** A residual U-Net is used to segment instances of cells using the fluorescence images of their H2B-GFP labelled nuclei. The U-Net consists of residual blocks and residual skip connections. A final 1⨯1 convolution layer, followed by softmax activation generates the output. Detected cells are localised using the centre of mass. **(B)** Cell centroids are used to crop nucleus-centred image patches from both transmission and fluorescence images, which serve as inputs to a CNN-based classifier to label instantaneous cell state. Labels indicate whether the cell is in interphase, mitosis or apoptotic. Scale bar = 10μm. (**C)** Using the localised and labelled cell observations, *btrack* uses Kalman filters to predict the future state of the cell 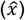 from previous observations. Each object appearing in the new frame has uniform prior probability of track association or loss. The belief matrix is an N x (M+1) matrix where N equals number of active tracks and M is the number of detected objects per field of view. Bayesian updates are performed using the predicted track positions and state information to calculate the probability that tracks are considered lost, or to be linked to an object. Objects which are not assigned, initialize new tracks. T1 represents a simple association, where the prediction matches the observation O1. In T2 the cell divides, meaning that there is no simple association. New tracks T3 and T4 are initialized. **(D)** An n-dimensional local search creates all possible hypotheses, each with an associated likelihood, for each track based on user defined parameters. Several hypotheses are generated for Track T2, as in **C**: possible linkage to T3 or T4, T2 undergoes mitosis generating T3 and T4, T2 is a false positive track or that T2 undergoes apoptosis. The likelihood of each hypothesis (P(E|h)) is calculated using the image and motion features. These hypotheses are evaluated in a global optimization step to generate the final tracking results.

In the next, optional step, the state of all identified cells was determined using an image classification network. The fluorescence images together with associated brightfield images are cropped to a bounding box centred on each cell and capturing the extent of the cell. Our convolutional neural network (CNN) classifier uses learnt image features to label each cell with one of the five states: interphase, prophase/prometaphase, metaphase, anaphase/telophase or apoptosis, with high accuracy (**Figure 2B, Supplementary information**).

Next, the detected cells are supplied to the tracking algorithm, which first assembles tracklets by linking cell detections over time that do not contain cell division events. We use a Kalman filter^33^, as a probabilistic model of each detected cell’s motion. The Kalman filter is used to learn and predict future positions, and their error, for all cells in the field of view (FOV). These predicted positions need to be matched with observations. We utilise a Bayesian approach for this data association step (**Figure 2C**). The Bayesian belief matrix approach enables incorporation of multiple probabilistic models, encompassing appearance as well as motion properties, into the early steps of the tracking algorithm. In the update stage, the posterior probability of each potential linkage is evaluated by constructing a belief matrix for all possible linkages. The belief matrix summarises the probability of each active track being assigned to each new object, and also the probability of that track becoming lost^34^. Further, by utilizing information from the cell state labelling, we can incorporate penalties for certain unwanted transitions (such as directly linking a child cell to a parent cell) during the update step. This improves the subsequent detection of mitoses, by ensuring that daughter cells are not incorrectly linked to parent cells.

However, for very large numbers of cells per FOV, the number of Bayesian updates required to track the cells can become very large. Therefore, we provide an additional method which performs approximate Bayesian updates during tracking, enabling rapid and efficient tracking of tens of thousands of cells. This approximation method limits the updates to a user defined search radius around each cell, and significantly improves performance, at the cost of assessing all possible track linkages. In all subsequent results, we utilize the exact, rather than approximate, computation method.

After linking observations into tracklets, they are next assembled into tracks and lineage trees using multiple hypothesis testing and integer programming^35,36^ to identify a globally optimal solution. We built an efficient hypothesis engine, which proposes possible origin and termination fates for each track based on their appearance and motion features (**Figure 2D**). The following hypotheses are generated: false positive track, initializing at the beginning of the movie or near the edge of the FOV, termination at the end of the movie or near the edge of the FOV, a merge between two tracklets, a division event, or an apoptotic event. In addition, we added ‘lazy’ initialization and termination hypotheses, allowing cells to initialize or terminate anywhere in the movie. These new hypotheses are strongly penalized, but their inclusion significantly improves the output by relaxing the constraints on the optimization problem. The log likelihood of each hypothesis was calculated for some or all of the tracklets based on the heuristics encoded in the hypothesis engine. The global solution identifies an optimal sequence of hypotheses that accounts for all tracklets. Once the optimal solution has been identified, tracklet merging can be performed. A graph-based search is used to then assemble the tracks into lineage trees, and propagate lineage information such as generational depth, parent and root IDs.

The final output consists of a binary mask of the segmented cells, centroids of each identified cell, cell state labels, trajectories, fate and a fully annotated lineage of each cell in the FOV. Once the image data are processed, *btrack* takes only a few seconds to completely track and assemble lineage trees, allowing quick feedback to the user. This compares to about 1.5-2 weeks of dedicated time^19^ of an experienced annotator to manually follow single cell trajectories over intermitotic time and organise the hierarchical relationships between ancestor and descendant cells into graphical tree representation, based on reconstruction of the ground truth trees for this publication. We provide a rigorous assessment of the segmentation, classification and tracking metrics in the supplementary information, and now focus on the retrieval of intermitotic times and lineage tree analysis.

### Benchmarking unsupervised lineage tree reconstruction

Reconstructing lineages represents a complex challenge and a common source of tracking errors in multigenerational cell observations. From our track validation dataset, we automatically reconstructed 154 lineage trees which contained at least one mitotic event. These trees were initiated in three instances: i) through their initial presence in the FOV at the beginning of a movie, ii) as a consequence of migration into the FOV, or iii) upon breakage of an existing branch from its tree.

Branch breakage occurred due to loss of a cell’s identity because of incorrect cell segmentation (e.g. due to low fluorescence signal) and was very rare (25 instances out of 728 cells organised into lineage trees, corresponding to one breakage per ∼6000 correctly linked frame observations). When the lost cell is re-detected, it becomes the founder of a newly initialized track. This track now becomes a root of a new ‘subtree’, which can continue to further divide or exist without further splitting (here referred to as ‘branch’).

To measure the correctness of parent-child relationships, we calculated the mitotic branching correctness score (MBC, **Methods**, a metric modified from previous studies^36^) in a lineage tree set of 24 randomly selected founder cells (**Supplementary information**). We scored the detection of mitosis as correct only when cell divisions were detected within a strict time window (± 1 frame; 4 mins) in computer-generated tracks relative to human-annotated ground truth lineage trees. Further, marking of false splitting events (e.g. two cells existing in close proximity at high cell density or misdetection of two cells in areas occupied by a single, large cell) could lead to erroneous generational relationships. To calculate the fidelity of generational depth, we introduced a penalty for excessive branching assignment. The progeny were considered as correct only when the generational depth relative to the tree root matched the ground truth.

In our testing tree pool comprising 24 lineage trees with 351 human-annotated cell division events, the MBC score for our *btrack* algorithm in both penalised and penalty-free variants, was 86.76% and 87.21%, respectively. This is approximately 2- to 3-fold greater than the benchmarking standards without and with the introduced generational depth penalty, respectively (**Supplementary information**). Our scoring system reveals the importance of our optional cell state classification step, which enables improved division hypothesis creation. Most other tracking methods employ hard distance thresholding to account for splitting events and, as a consequence, struggle to discriminate between over-segmentation artifacts, two independent cells migrating into close proximity, and true mitotic events.

The correctness of mitotic branching is directly reflected in the leaf retrieval score (LRS, **Methods**), which computes the number of all terminal cells (cells with no identified children) in the respective lineage tree appearing in the last frames of the movie, with the correct generational depth relative to their founder cell. Keeping record of cell division history over the entire duration of the live-cell imaging is necessary to elucidate whether cells with shorter cell cycles have the capacity to dominate the population and competitively outgrow the slowly dividing cells over time.

Here, we report an LRS of 84% on 364 leaf cells in the ground truth lineages, almost 3-fold higher than for other benchmarked trackers (**Supplementary information**). For ease of lineage tree visualisation, we developed a dockable widget for real-time feedback between tracking data and source images based on napari visualisation software^37^. Within the GUI, we provide a convenient visualization for track survival over time (**Supplementary information, Supplementary Video S1**), revealing trajectories that can be tracked to the movie start via their ancestors, as an easy way to assess tracking fidelity and identify errors.

Finally, we computed metrics reflecting how well the human-annotated cell trajectories are followed by the reconstructed lineage trees (recall, **Methods**) and vice versa (precision, **Methods**)^36^. Without any manual curation, our *btrack* framework faithfully captured 90.6% of ground truth track observations (recall) with over 99.7% of observations in agreement with the ground truth trees (precision). Taking further advantage of our large dataset, we randomly sampled an additional 302 unseen, automatically reconstructed trees from multiple movies. For each tree, we visually scored them as sufficient if they had >90% recall of the true tree as determined by an expert annotator. Overall, 201 out of 370 reconstructed trees had >90% recall. The remainder of the trees contained one or more errors. To quantify these, we returned to our fully annotated test dataset. In the test dataset, we needed to perform only eight ‘subtree’ re-assembly actions to the original trees (**Supplementary information**), which added 11,505 frame observations, and a single subtree swap (where part of the tree was falsely associated with another tree). These re-assemblies increased the recall to 98.9%. Further ‘branch’ stitching, requiring additional 24 user-mediated curations, adding another 1,926 frame observations, increased recall to 99.8% of the full human-annotated trees. However, the value of highly effective tracking may come at the expense of the user’s time to curate tracks and may not significantly impact the output. These results show that our *btrack* tracking algorithm is able to reconstruct cell trajectories with excellent performance in lineage tree reconstruction, even without manual curation.

### Single cells exhibit substantial heterogeneity in cycling durations and colony expansion

Having validated the lineage tree reconstructions, we pooled the tracking data from the entire dataset of 44 time-lapse movies. First, we filtered these cells to remove any with unknown, partially resolved parents (root cells) or progeny (leaf cells). This yielded 22,519 cells, organised into lineage trees spanning up to eight generations. Next, we calculated the per-cell intermitotic time as the time between the first appearance of separated chromosomes during mitosis (labelled as anaphase by the CNN; due to the temporal sampling this may be before cytokinesis occurs) to the frame preceding the next anaphase (**Figure 3A**).

**Figure 3.**
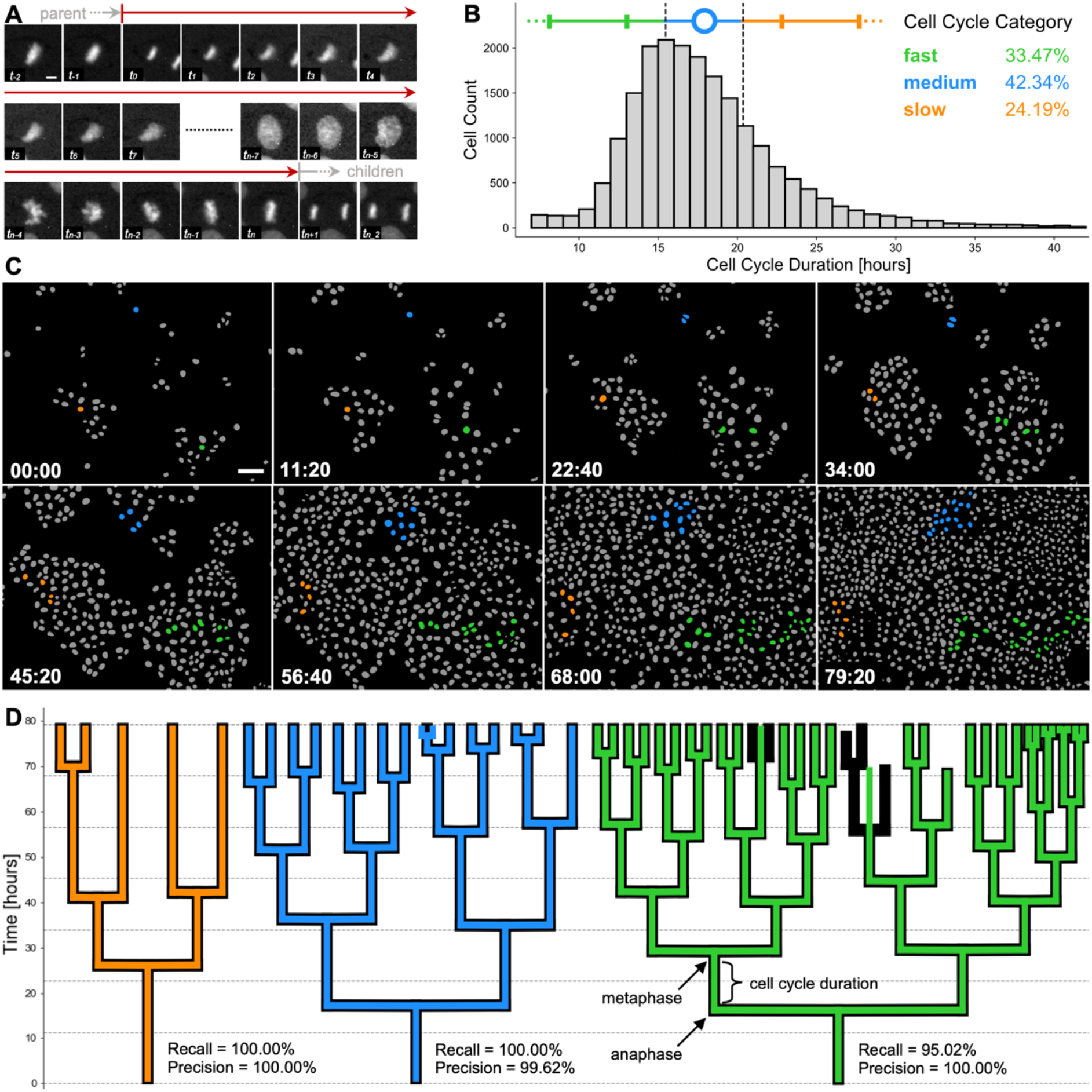
Cell cycling heterogeneity and colony expansion capacity within single cell clones. **(A)** Cell cycle duration, defined here as the time elapsed between two subsequent anaphases, from the mitosis of reference cell’s parent (first two images) to division into two children cells (last two images). Scale bar = 5 um. **(B)** Calculated cell cycle duration distribution over the pooled MDCK live-cell imaging dataset (n=20,074 single cells from 5,325 unique lineage trees). Error bars (solid vertical bars) show one and two standard deviations around the mean (blue circle) of tracked cells for generational depths spanning 1 to 6. Cell categorisation into fast (green), medium (blue) and slow (orange) dividers is depicted as below, within or above 1/2 standard deviation away from the mean (dashed vertical lines), respectively. Percentages of single cells belonging to each category are stated. **(C)** Sequence of 8 colourised binary masks with segmented individual cells (grey) on background (black), highlighting cell proliferation from the start (top left) to the end (bottom right) of a representative movie. Time in hh:mm is indicated in the bottom left corner. Founder cells and progeny corresponding to slow (orange), medium (blue) and fast (green) dividers are highlighted. Scale bar = 50μm. **(D)** 2D representations of typical lineage trees captured from data, showing slow (orange), medium (blue) and fast (green) dividing cells. Computer-generated tracks (colour) are overlaid on human-reconstructed ground truth trajectories (black). Individual recall and precision scores are shown for each lineage tree, with vertical branch lengths corresponding to intermitotic time (cell cycle duration) elapsed between the first post-division frame (anaphase) to the last frame prior to the next cell division (metaphase). Trees illustrate an error-free tracking (orange tree) and 2 types of tracking errors, i.e. falsely identified mitosis (blue tree) and missed mitotic event (green tree).

Detailed inspection of the nucleus size and CNN labels were utilised to determine bounds for cell exclusion from our distribution (**Supplementary Figure S3**). We found that the tracks with intermitotic time below 7 hours had high incidence of start with non-anaphase label, did not end with pro-(meta)phase label, or a combination of both. This observation suggested that these short tracks represent fragments of cell trajectories where branch breakages occurred, rather than being representative of ultra-fast cycling cells. On the other hand, visual observation of the nuclear growth (increase of cell nucleus segmentation mask area over time) indicated that tracks which were calculated to have division time longer than 42 hours captured track instances where a parent cell (undergoing mitosis) was falsely linked to one of the arising children cells, most likely due to imperfection in the segmentation step.

Therefore, to avoid possible incorporation of prematurely terminated tracks, concatenated parent-to-child tracks or other tracking errors, we filtered our pooled dataset to only consider cells with cycling lengths between and including 7 and 42 hours for further analysis (**Figure 3B**). Our final dataset consisted on 20,074 cells with known lineage over up to 6 generations, representing at least two orders of magnitude greater numbers than in previous studies^38,39^. Importantly, the whole process to filter the relevant cells, calculate their division times and pool the single-cell information across 44 movies took less than a minute to produce, which greatly favours our tracking pipeline for large data analyses.

The pooled dataset shows a positively skewed normal distribution of cell cycle durations (mean ± standard deviation, 17.9 ± 4.9 hours, **Figure 3B**). This distribution confirms the presence of division time heterogeneity at very large population numbers, while standing in good agreement with previously published work^38–40^. We also confirmed that cell cycle lengths showed no trend over the duration of live-cell imaging with respect to cell birth time relative to start time for the first 60 hours of time-lapse imaging (**Supplementary Figure S8**).

We used the population distribution to categorise individual cells as fast, medium and slow dividers based on whether their cycling duration was below (15.5 hours), within or above (20.3 hours) half a standard deviation away from the sample mean (**Figure 3B**). This definition yields a subpopulation of fast dividing cells (n=6,719 cells; 33.5% of the population), medium dividers (n=8,500 cells; 42.3%) and slow dividers (n=4,855 cells; 24.2%).

We extended this categorisation to describe individual cell families and contrasted three lineages extracted from our representative movie, with the trees observed from the movie start frame. Mapping the number of progeny originating from these three founder cells (**Figure 3C, Supplementary Video S2**), we confirmed our large-scale observation of broad cycling heterogeneity spectrum. Based on the average cell cycle length of all fully-resolved cells within the tree, we classified the cell families as slow, medium and fast cyclers. Visual inspection of their lineage tree representations reveals that variability in intermitotic durations amongst single cells directly influences the cell capacity to divide and poses potential for fast cycling cells to eventually dominate the population by overgrowing the slower-cycling clones (**Figure 3C-D**), as suggested previously^5^.

Indeed, taking trees corresponding to three different root cells, we find that at the end of the movie, the slow dividing family results in 5 leaf cells, the medium dividing family in 15 cells and the fast dividing family in 31 leaf cells, over the same period of 80 hours. These correspond to mean intermitotic durations of 20.6 ± 6.5 hours (n=3 cells), 16.9 ± 4.5 hours (n=14 cells), and 14.0 ± 1.9 hours (n=29 cells) for each tree, respectively. Our findings suggest a high degree of intrinsic cell cycling heterogeneity present in the wild-type MDCK cell population. This cycle-time heterogeneity appears to be maintained across cell lineages.

### Deep lineage trees enable evaluation of correlations of cell cycle durations across generations

Next, we studied whether the observed cell cycle duration is inherited upon cell division. Cell cycle duration may be governed by stochastic processes, or inherited upon cell division in a deterministic fashion which only appears to be random, as posed by others^4^. Previous studies suggested that cell cycle durations within lineages show poor correlation when observing the direct ancestral cell pairs (mother-daughter and grandmother-granddaughter), but remain highly correlated when examining intra-generational relationships (sister and cousin cells)^4,7,8^. However, due to lack of deep datasets, the analysis has traditionally focussed on immediate cell relatives, including mother, grandmother, sister and first cousin relationships^4,7^ and less frequently on more remote family members, such as great-grandmothers or second cousins^8^. We have extended previous studies using the enhanced depth (up to eight generations), breadth (up to fourth cousin distance from reference cell), and number of lineages (5,032 trees) from our automatically reconstructed lineage tree pool (**Figure 4A**). If the cell cycle duration is not controlled in a heritable fashion, we would expect poorly correlated cycling durations across ancestral as well as side-branch family relationships. This would, however, be in contrast with the strong deterministic control which would likely yield highly correlated family relationships maintained across several generational depths as a result of heritable biochemical and/or mechanical factors.

**Figure 4.**
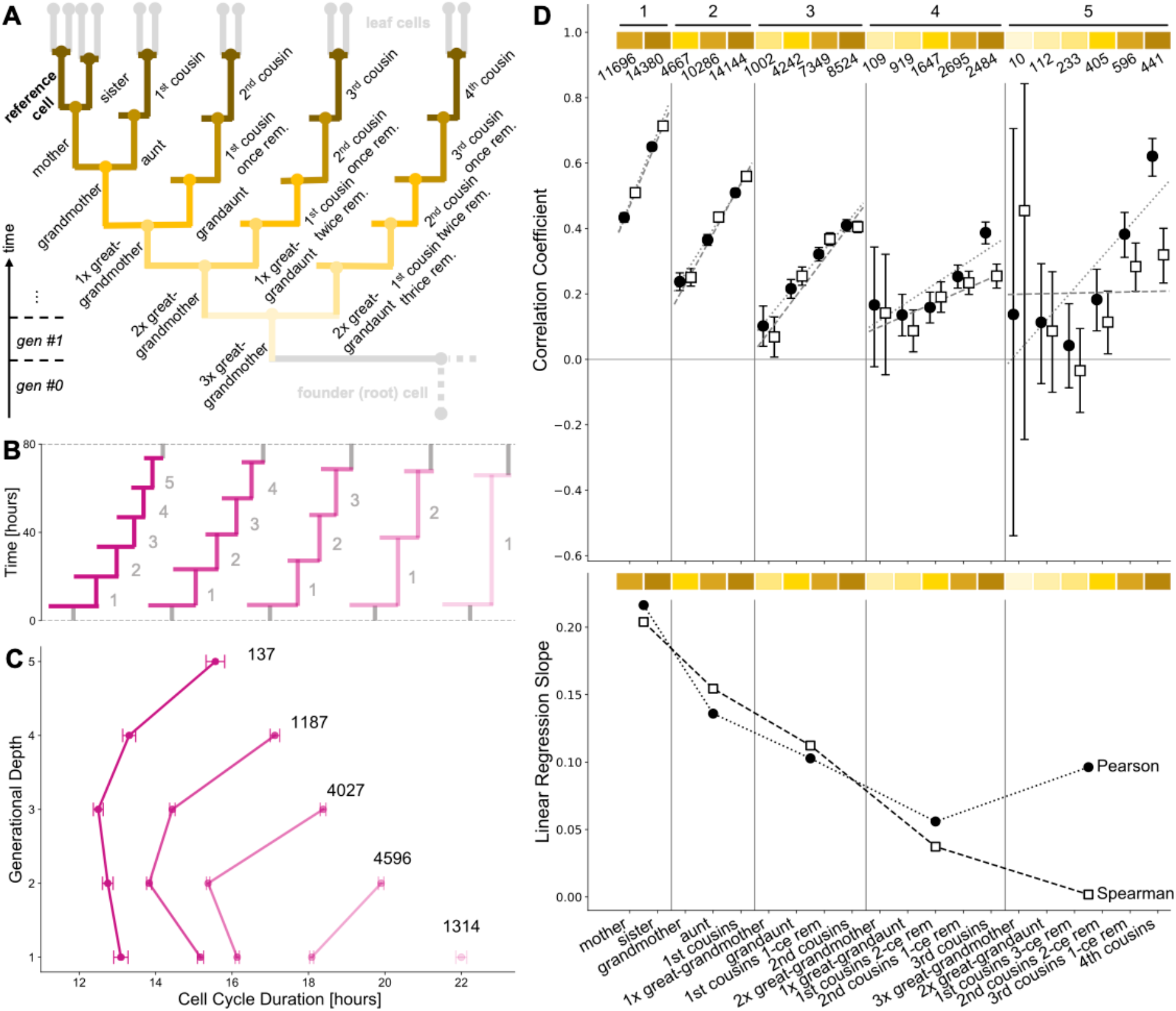
Large-scale multigenerational analysis of single-cell cycling durations. **(A)** Illustrative lineage tree showing 20 types of family relationships of a reference cell (bold) to its lineage relatives determined using our automated approach. The tree captures extended kinships to great-great-great-grandmother distance (directly ancestral) to fourth cousin span (generationally equal). Strength of the branch colour shading illustrates the generational distance to nearest common ancestor between family member and reference cell. Generational depth assignment with respective to the tree founder cell (bottom left) is illustrated. **(B-C)** Cell cycling speed influences the amount of populated progeny in lineage trees. Fully-resolved lineage paths categorised according to the number of completed divisions, from 5 (leftmost, fastest dividers) to 1 (rightmost, slowest dividers). Plotted is the average cell cycle division time for each generational depth in the lineage path. The cell subpopulation with shorter average intermitotic times completes more division cycles than the slower-cycling cells. The number of technical replicates is shown. Error bars indicate standard error of the mean. **(D)** *(Top)* Pearson (black circle points, dotted line) and Spearman (white square points, dashed line) rank correlations of cycle lengths between relatives from 5,032 unique lineage trees. Linear regression of each group is shown. Error bars indicate 95% confidence interval. Shaded golden blocks relate to the family relationships as in A, with counts of analysed cell replicates per kinship type listed. Horizontal black bars connect family relatives with equal generational distance to the nearest common ancestor with respect to reference cell, as indicated by numbers above the bars. *(Bottom)* Linear regression slopes for both coefficient types linearly decrease for at least 4 generational distances to nearest common ancestor.

First, we examined 11,261 lineage paths in our dataset. These lineage paths are determined for cells with an unbroken lineage from the founder cell, tracked across the entire movie duration. We calculated lineage paths originating from 2,133 unique lineage trees, and divided these paths according to their depth, *i*.*e*. the number of intermediate cells present in the path which were neither root nor leaf (**Figure 4B**). For each path, we computed the mean cell cycle duration per generational depth. Without any further categorisation, the average cell cycle durations cluster together according to their path depth (**Figure 4C**). As expected, the cell cycle duration was shown to be indirectly proportional to the number of divisions the cell completes within its lineage path, with cells in 5-member paths having the shortest-cycling times across all generational depth as compared the longer-cycling cells. Our analysis also revealed that the cell cycle duration is elongated in later generations (**Figure 4C**). This behaviour may be due to contact inhibition arising from cell crowding or to nutrient depletion, as proposed by others^38^.

Next, we extracted 20 different types of cell pair relatedness (**Figure 4A**) and calculated the correlation between cell cycle duration in these pairs. We computed the cell cycling correlations in family member pairs from the trees with direct ancestral (mother, grandmother, great-grandmother, etc.) or side-branch lineage relationships at identical (sister, first cousins, second cousins, etc.) of different (aunt, grandaunt, cousins once removed, etc.) generational depth compared to the reference cell (**Figure 4A**). In doing so, we were able to analyse orders of magnitude greater numbers and significantly greater depth of lineage trees then in previous studies^4,7,8^. Our ancestral analysis comprised 11,696 mother, 4,667 grandmother, 1,002 great-grandmother, 109 great-great-grandmother and 10 great-great-great-grandmother comparisons **(Figure 4)**.

We observed moderate correlations (Pearson and Spearman rank coefficients of 0.43 and 0.51, respectively) between 11,696 reference and mother cell pairs (**Figure 4D**), which represents an order of magnitude higher, yet still relatively low, mother-daughter correlation compared to previously shown trends^4,7,8^. The correlations continued to decrease with increasing ancestor distance all the way to a cell’s great-great-grandmother (0.24, 0.10, 0.17 for Pearson and 0.25, 0.07, 0.14 for Spearman rank coefficients). Such diminishing ancestral correlations would be strongly indicative of cell cycle duration inheritance being a stochastic event, as previously speculated^4^.

In contrast, we observe relatively conserved correlations in progressively remote side-branches capturing cell pairs with identical generational depth. Indeed, sister cell cycling times were highly correlated (Pearson and Spearman rank coefficients of 0.65 and 0.71, respectively, 14,380 sister cells examined). Further, cell cycle times remain moderately correlated across the lineages all the way to the fourth cousin distance (**Figure 4**). The transition from poorly correlated direct ancestors to relatively well correlated generationally equal family side-branches is characterised by increasing correlations of cells with varying kinship similarity and lineage proximity to the reference cell (**Figure 4A**), including aunts (10,286 comparisons) to reach first cousins, grandaunts (4,242 comparisons) and cousins once removed (7,349 comparisons) to access second cousins, etc. The steepness of the linear regression slope calculated from the respective correlation coefficients per each branch was shown to decrease with increasing generational distance to first common ancestor shared with the reference cell (**Figure 4**). Our results derived from high-replicate lineage data are in good agreement with previously published studies, using manually annotated data^4,7,8^. Overall, the data confirm that cycling length is correlated across longer-range relationships (4 and 5 generations to nearest common ancestor) than previously examined. These long-term correlations may suggest heritability in cell cycle durations, as proposed by others^5^.

## DISCUSSION

We developed an easy-to-use, open source Python package to enable rapid and accurate reconstruction of multi-generational lineage trees without time-consuming manual curation. This allows users to characterise population-level relationships with single-cell resolution from time-lapse microscopy data.

As an illustration, we used our software to analyse heritability and stochasticity in cell cycle durations in isogenic populations of epithelial MDCK wild-type cells. We analysed: (i) cell cycling heritability in individual clones, (ii) cell doubling duration heterogeneity within populations and (iii) cycle-length correlations in extended cell families. We successfully recovered the experimental trends from previous studies that used manual or semi-manual annotation^4,7,8,38,39^, and extended the analysis with orders of magnitude more experimental data, enabled by our fully automated approach. Further, as a result of our approach, we were able to extend the cell kinship correlation analysis in both depth and breadth of studied lineage relationships, which was not possible in previous work due to lack of experimental data. Our data suggest that heritable cell cycling is present, the regulation of which may arise from tightly controlled, deterministic processes, as proposed by others^5^. Surprisingly, cell cycle times are tightly distributed within lineages in our dataset. It is possible that, as culture conditions evolve with time (nutrients become depleted and cells more confluent), the highly correlated behaviour between same generation family members may be a consequence of environmental synchronisation. Further experiments accounting for density and nutrient depletion are necessary to distinguish these hypotheses. Our approach, using high-replicate lineage data will enable study of the factors influencing the accumulation of noise in correlations with respect to lineage distance by providing an accurate, rapid, and easy to use analysis package.

In summary, our study demonstrates the utility of fully automated deep lineage tree reconstruction from time-lapse microscopy data. We developed a powerful software package to enable users to analyse cell cycle durations at the single-cell level over large cell populations and explore the relationships of cells assembled into lineage trees spanning up to eight generations in an unsupervised manner. The potential applications of this approach include, but are not limited to, analysis of (cancer) stem cell identification in tissues, detection of differentiation and/or reprogramming success in populations, studying of cell cycle control mechanisms in cell lineages and dynamic high-throughput screens of various pharmaceutical compounds with live-cell imaging.

## METHODS & MATERIALS

### Automated widefield microscopy

A custom-built automated epifluorescence microscope was built inside a standard CO_2_ incubator (Hereaus BL20) which maintained the environment at 37^°^C and 5% CO_2_. The microscope utilised an 20x air objective (Olympus Plan Fluorite, 0.5 NA, 2.1mm WD), high performance encoded motorized XY and focus motor stages (Prior H117E2IX, FB203E and ProScan III controller) and a 9.1MP CCD camera (Point Grey GS3-U3-91S6M). Brightfield illumination was provided by a fibre-coupled green LED (Thorlabs, 530nm). GFP and mCherry/RFP fluorescence excitation was provided by a LED light engine (Bluebox Optics niji). Cameras and light sources were synchronised using TTL pulses from an external D/A converter (Data Translation DT9834). Sample humidity was maintained using a custom-built chamber humidifier. The microscope was controlled by MicroManager^41^ and custom-written software OctopusLite.

### Cell Culture and Time-Lapse Movie Acquisition

We used Madin-Darby Canine Kidney (MDCK) epithelial cells as a model system. Wild-type MDCK cells were grown, plated and imaged as described previously^40,42^. Briefly, MDCK cells were transduced using Lentivirus to stably express fluorescently tagged histone marker to enable visualisation of nucleic acid organisation during cell cycle. The established MDCK cell line expressing H2B-GFP fusion protein was seeded at initial density of approx. 3 ⨯ 10^4^ cells / cm^2^ in 24-well plates (ibidi). Imaging was started 2–3 h after seeding. Imaging medium used during the assay was phenol red free DMEM (Thermo Fisher Scientific, 31053) supplemented with tetracycline-free bovine serum (Thermo Fisher Scientific, 10270106), and antibiotics. A typical experiment captured multiple locations with a field of view of approx. 530 ⨯ 400 μm for over 80 hours with constant frame acquisition frequency of 4 minutes for each position. Multi-location imaging was performed inside the incubator-scope for duration of approx. 1200 images (80 hours).

### Image Processing

We constructed a 2D U-Net^32^ for segmenting cells in the time-lapse microscopy images. Our U-Net architecture consisted of four down layers and four up layers, with convolutional and interpolated upscaling transformations, respectively (**Figure 2A**). Rather than feature concatenation in the skip connections, we used element-wise addition to resemble a Residual Network approach (ResNet^43^). Further, we used residual blocks in the convolutional blocks to minimize vanishing gradients during training. The calculated pixel-wise weight maps forced the network to prioritize regions separating proximal nuclei. We hand-segmented 150 images to provide ground truth images for network training and validation. During training, regions of 768⨯768 pixels were randomly cropped from the source images, using on-the-fly augmentations such as cropping, rotation, flipping, noise addition, uneven illumination simulation, scale and affine deformations. We trained the network using the Adam optimizer^44^ with batch normalization, a batch size of 16 for 500 epochs. We assessed the performance of the network using a held-out test set dataset (comprising 20 images), calculating intersection over union (IoU) and other matching scores (**Supplementary Information**).

Next, we localized each cell by the centre of mass of each segmented region, and used a convolutional neural network (CNN, **Figure 2B**) classifier^40^ to label the state of each cell. We performed the CNN training with >15,000 manually annotated fluorescent & transmission examples and assessed network performance calculating the confusion matrix with a set of 2500 validation images, which were not used for training (**Supplementary Information**). We used the same augmentation procedure as with the U-Net and trained with a batch size of 128 using the Adam optimizer. Segmented and classified cells were written out in HDF5 format files. Cell tracking was performed as described. All image processing was performed in Python, using scikit-image, scikit-learn, Tensorflow and Keras libraries, on a rack server running Ubuntu 18.04LTS with 256 Gb RAM and NVIDIA GTX1080Ti GPUs.

### Validation of lineage tree reconstruction

We use the following metrics to validate the lineage tree reconstruction. The Mitotic Branching Correctness (MBC), a measure of the ability of the tracker to identify mitoses, is calculated as:

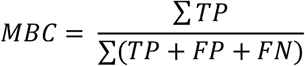

Where true positive (TP) mitotic events are described as track splitting events identified within time distance θ_t_ of the ground truth (GT) (**Supplementary Figure S4**). We use a strict threshold of θ_t_ = ±1 frame. To consider the mitotic event as a TP, both the parent (single dividing track) and progeny (two newly appearing tracks) cells must exist in a correct generational depth relative to the tree root (founder cell) (**Supplementary Figure S5**). False positive (FP) mitoses were calculated as the difference between the total events detected by the tracking algorithm minus total TP events. False negative (FN) mitoses represent the difference between ground truth mitoses count and total TP events. In addition, we calculate the Leaf Retrieval Score (LRS), a measure of the number of correctly recovered tracks at the end of the movie, is calculated as:

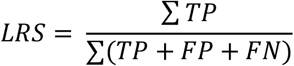

Where leaves (a terminal cell of a tree which doesn’t further divide) are considered as TP when its entire lifetime matched the GT observation. Additionally, an exclusion of all leaves which were not followed until the last movie frame (**Supplementary Video S1**) was introduced to only include TP leaves as those appearing at the movie end. FP leaves were calculated as the difference between the total leaves detected by the tracking algorithm minus total TP leaves. FN leaves represent the difference between the ground truth leaf count and total TP leaves.

The recall (also known as Target Effectiveness), is essentially the ability of the tracker to correctly recall trajectories present within a lineage tree, and is calculated as:

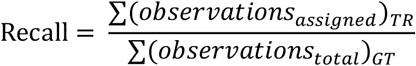

Where the number of assigned track observations of the target is divided by the overall number of frames in ground truth target. Finally, we calculate the precision (also known as Track Purity), which represents the ability of the tracker to reconstruct the trajectories present within the lineage tree, as is calculated as:

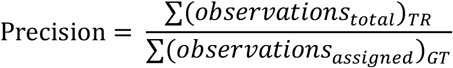

Where number of total ground truth tracked frames is divided by the overall number of assigned track observations followed by the tracker. The track purity score is designed to account for FP events and expresses the precision of the tracker.

### Software Implementation & Data Availability

The *btrack package* was implemented in Python 3.7 and C/C++ using CVXOPT, GLPK, Numpy and Scipy libraries. We tested the software on OS X, Ubuntu and Windows 10. In addition, we developed interactive track and lineage tree visualization software named *Arboretum. Arboretum is* developed as an extension for the open-source multi-dimensional image viewer, *napari*^37^. We have provided extensive documentation online, including supplementary user-friendly tutorials with tracking examples, example data and installation instructions. Links to the source code and example datasets are available at: https://github.com/quantumjot/CellTracking.

## ACKNOWLEDGEMENTS

Funding. KU is supported by a BBSRC LiDo studentship. GV is supported by BBSRC grant BB/S009329/1 to AL and GC.

## Author Contributions

GC and AL conceived and designed the research. KU and GV performed experiments. KU developed and performed computational analysis. AL wrote the image processing and cell tracking code. KU, GV, GC, and AL evaluated the results and wrote the paper.

We thank members of the Lowe and Charras labs for helpful discussions.

