## Supplementary Information for "Automated deep lineage tree analysis using a Bayesian single cell tracking approach"

*Tracking Software Calibration:*

One of the major challenges of cell tracking is the propagation of errors throughout the pipeline. Each step gives rise to the possibility of incorporating errors such as premature track breakage, misidentification of cell splitting events and incorrect assignment of parent-children relationships upon lineage reconstruction. Because accumulation of small unaddressed errors over many time points can lead to a substantial drop in tracking accuracy, we performed a rigorous assessment to test our pipeline performance.

We define 4 different types of observations in the tracking output:

- *true positive* (TP, hit), which represents a true detection where the same object is found in the output and manual annotation
- *false positive* (FP, ghost), which represents a detection in the computer-generated output that does not appear in the manual annotation
- *false negative* (FN, miss), which represents an undetected object in the computer-generated output which is present in the manual annotation
- *identity swap* (IS, mismatch), which represents a tracking-related error where (i) the unique identity of two tracked trajectories are swapped, most likely due to close proximity or with crossing trajectories, (ii) the incorrect linkage event leads to assignment of a new ID label to an existing track, or (iii) the misidentification of mitosis, where one of the children cells becomes falsely concatenated to the parent cell.

**Equation 1 | Multi-Object Tracking Precision (MOTP)**

$$MOTP = \frac{\sum \sqrt{(x_{GT} - x_{TR})^2 + (y_{GT} - y_{TR})^2}}{\sum (objects_{TP})}$$

Where the Euclidean distance between centroid *(x,y)* coordinates of manual annotations (GT) and tracker outputs (TR) is computed and divided by sum of all within distance threshold $\theta_d$ of 20 px away from ground truth (GT) observations at time *t*. This metric represents how precisely tracking algorithm can determine the position of an object. It is the ratio of the total error in position to the number of true positive correspondences between the tracker and manual annotation.

**Equation 2 | Intersection over Union (IoU)**

$$IoU = \frac{TP}{TP + FP + FN}$$

Where the ratio between common shared pixels (true positive, TP) and sum of pixels of two segmented objects, including the shared pixels (true positive, TP, false positive, FP, false negative, FN) is computed per single object between the observations and the ground truth.

**Equation 3 | Multi-Object Tracking Accuracy (MOTA)**

$$MOTA = 1 - \frac{\sum_t (FN_t + FP_t + IS_t)}{\sum_t (objects_{GT})_t}$$

Where sum of all trajectory observations subjected to tracking errors (FN, FP and IS) at short time-lapse sequences of length *t* is divided by the total number of observations in the ground truth (GT) at time series *t*. Time series *t* = ~20 frames. This metric represents the errors associated with detecting objects and accurately keeping track of them, independent of the ability to precisely localise them.

*Assessment of tracking metrics:*

We selected representative time-lapse microscopy movies, and for each, shortlisted 3 representative frames capturing cells grown to low, medium and high confluency and manually annotated the nuclear areas to calculate the fidelity of cell detection and localisation. We then manually reconstructed the ground truth lineage trees of 24 founder cells (representing more than ⅓ of initially seeded cells), the subsequent progeny of which (1,032 cells including tree founders) spanned the entire movie duration. The performance of our movie analysis workflow was contrasted to a Python-based package, *TrackPy*[1], and an ImageJ/FiJi plugin, *TrackMate*[2]. Both of these cell detection and tracking engines represent recently-developed popular particle tracking frameworks which function as backbones to the current state-of-the-art cell tracking software, such as *Usiigaci*[3] and *MaMuT*[4], respectively.

To assess the quality of the cell detection step, we measured the multi-object tracking precision (MOTP, **Eq 1**) in cells appearing in three manually labelled frames capturing fluorescently labelled cell nuclei sampled at low, medium and high confluency. Out of 869 cells in total, 847 cells had their centroid coordinates estimated within threshold of 20 pixels from designated cell in ground truth. Our pipeline localised the cells with highest precision (mean error 0.99 pixels), performing with at least 2.5-fold lower localization error than other benchmarked cell detectors (**Figure S1A**).

Unlike with other benchmarked pipelines, which output only the nucleus centroid coordinates, our localisation method performs a pixel-wise image classification, which offers the opportunity to segment the whole area belonging to the cell nucleus from which centroid coordinates are calculated. To calculate the accuracy of the nucleus area segmentation, we computed the per-object intersection over union metric (IoU, **Eq 2**) of the areas for each individual nucleus from the U-Net and how they compare to the manually labelled ground truth areas. We introduced a strict IoU thresholding (0.5 to 0.9 with 0.1 increments), i.e. we only scored a nucleus as hits (true positive, TP) when the computer-generated mask overlapped with the ground truth label by at least 50% (IoU 0.5), and above. We report that out of 616 ground truth nuclei in a fully confluent FoV (**Figure S1B**), 574 nuclei were sharing areas with at least 50% overlap, yielding a Jaccard Index of 0.933 (93% of objects detected in the FoV). As expected, the calculated localisation error was decreasing as only the highest overlapping cells are included in calculation of progressively increasing IoU threshold values. More details can be found at: https://github.com/quantumjot/unet_segmentation_metrics.

Next, we calculated the multi-object tracking accuracy (MOTA, **Eq 3**) which scores the tracker's ability to retain cell's identity and trajectory over longer periods of time. MOTA score intrinsically penalises the tracking pipeline for static (ghost or missed objects) as well as dynamic (identity switches) errors, which often strongly rely on the detection algorithm performance. We used short movie sequences of up to 20 successive fields of view at three confluency levels as previously. Observing 2,161 individual cell objects over time, our tracking pipeline (97.66%) performed comparably to TrackMate (97.54%), but significantly outperforms the TrackPy (73.77%) tracking algorithm.

Localization accuracy, as determined by the different algorithms used, accounts for the majority of differences between the MOTA scores for different tracking packages. Importantly, our library is agnostic to the segmentation and localization methods used to identify the cells in the image data. The requirement for the subsequent tracking step is to process the raw sequence of input images into a vector of localisation coordinates of individual objects appearing in each field of view. Optionally, it is desirable to describe each object with a numerical label which corresponds to the cell state based on its progression towards mitosis (0 - interphase, 1 – pro-/metaphase, 2 - metaphase, 3 – ana-/telophase, 4 - apoptosis), which is an additional input to the tracking algorithm for improved performance. The simple structure of tracking data input enables users to choose, or design, a pipeline appropriate to their data and integrate the tracking algorithm directly into their image analysis pipeline.

We used a representative movie comprising 1193 frames with 1600 x 1200 px for cell detection by the tracker. The localisation blob diameter was optimised for the entire movie on 3 fields of view representative of 3 confluency levels (low at 14%, medium at 41% and high at 98%).

**TrackPy**

We selected the 23 px diameter estimate which yielded the most satisfactory results (66, 180, 574 detected objects vs. 68, 185, 616 objects found in ground truth). All objects below the 'minmass' of 500 were filtered to exclude ephemeral blobs from the analysis. For track linking, the maximum displacement was specified to be 25 px and the memory for missed detections was set for 3 frames. The tracking yielded 4,521 trajectories in total, out of which 1,010 trajectories were tracked between the range of 7-42 hours.

**TrackMate**

The representative movie had the 'z' and 't' dimensions swapped upon loading to TrackMate. The blob detection calibration settings remained unchanged as in default (pixel width, height and depth equal to 1.0 px, time interval of 1 frame). Downsample Laplacian of Gaussian (LoG) filter was used for detection, with the sigma suited to the blob estimated size. Estimated diameter was chosen to be 30.0 pixels with threshold set to 0.0 (default), and downsampling factor of 4.0. Initially, 1,302,106 blobs were identified throughout the movie, subjected to initial thresholding for quality of 0.73 to retain 377,291 detected objects. For subsequent tracking step, the Linear Assignment Problem (LAP) mathematical framework was selected. Parameter for maximal distance in frame-to-frame linking was set to 30.0 px, track segment gap closes option was allowed and set to maximum distance of 15.0 px and 2-frame gap. Track segment splitting was allowed below the 25 px threshold. No feature penalties or additional track filtering were introduced in the tracking pipeline. Due to large number of lineage trees in the tracking output, the attempt to visualise all of the lineage data at once using the in-built 'TrackScheme' GUI was unsuccessful. The results tables produced via 'Analysis' option were saved out and computationally processed outside of the TrackMate interface using a custom-written software to reconstruct trees from linked spots information (**Supplementary Figure S6**). The tracking yielded 2,684 trajectories in total, out of which 1,264 trajectories were tracked for the duration between 7-42 hours with no splitting events.

### Table S1 | Summary of the Datasets Used for Tracking Performance Analysis.

| | Biological Replicates | Technical Replicates | Lineage Tree Pool Size | Fully Resolved Single Cells |
|---|---|---|---|---|
| *Total Movies Included in Analysis* | 9 | 44 | 5,032 | 20,074 |

### Table S2 | Summary of the Cell Pairs Used for Family Tree Cycling Correlations.

| Kinship Type | Cell Pair Count | Pearson Rank | | | Spearman Rank | | |
|---|---|---|---|---|---|---|---|
| | | Correlation Coefficient | Lower 95% confidence bound | Upper 95% confidence bound | Correlation Coefficient | Lower 95% confidence bound | Upper 95% confidence bound |
| *mother* | 11696 | 0.42 | 0.42 | 0.45 | 0.51 | 0.5 | 0.52 |
| *sister* | 14380 | 0.65 | 0.64 | 0.66 | 0.71 | 0.71 | 0.72 |
| *grandmother* | 4667 | 0.24 | 0.21 | 0.26 | 0.25 | 0.22 | 0.28 |
| *aunt* | 10286 | 0.37 | 0.35 | 0.38 | 0.44 | 0.42 | 0.45 |
| *1st cousins* | 14144 | 0.51 | 0.5 | 0.52 | 0.56 | 0.55 | 0.57 |
| *1x great-grandmother* | 1002 | 0.1 | 0.04 | 0.16 | 0.07 | 0.01 | 0.13 |
| *grandaunt* | 4242 | 0.22 | 0.19 | 0.24 | 0.25 | 0.23 | 0.28 |
| *1st cousins 1-ce rem.* | 7349 | 0.32 | 0.3 | 0.34 | 0.37 | 0.35 | 0.39 |
| *2nd cousins* | 8524 | 0.41 | 0.39 | 0.43 | 0.41 | 0.39 | 0.42 |
| *2x great-grandmother* | 109 | 0.17 | -0.02 | 0.34 | 0.14 | -0.05 | 0.32 |
| *1x great-grandaunt* | 919 | 0.14 | 0.07 | 0.2 | 0.09 | 0.02 | 0.15 |
| *1st cousins 2-ce rem.* | 1647 | 0.16 | 0.11 | 0.21 | 0.19 | 0.14 | 0.24 |
| *2nd cousins 1-ce rem.* | 2695 | 0.25 | 0.22 | 0.29 | 0.23 | 0.2 | 0.27 |
| *3rd cousins* | 2484 | 0.39 | 0.35 | 0.42 | 0.25 | 0.22 | 0.29 |
| *3x great-grandmother* | 10 | 0.14 | -0.54 | 0.71 | 0.45 | -0.25 | 0.84 |
| *2x great-grandaunt* | 112 | 0.11 | -0.07 | 0.29 | 0.09 | -0.1 | 0.27 |
| *1st cousins 3-ce rem.* | 233 | 0.04 | -0.09 | 0.17 | -0.03 | -0.16 | 0.09 |
| *2nd cousins 2-ce rem.* | 405 | 0.18 | 0.09 | 0.28 | 0.11 | 0.02 | 0.21 |
| *3rd cousins 1-ce rem.* | 596 | 0.38 | 0.31 | 0.45 | 0.28 | 0.21 | 0.36 |
| *4th cousins* | 441 | 0.62 | 0.56 | 0.68 | 0.32 | 0.23 | 0.4 |


**SUPPLEMENTARY MOVIES:**

**Video S1 | Separation of automatically tracked single-cell lineages.** Colours code for 'survivor' cells which can be fully tracked to the movie start through their lineage (cyan), 'incomer' cells which migrated into the field of view throughout the duration of the imaging (yellow) and 'mistracked' lineages where a tracking error, such as trajectory breakage or falsely identified mitosis, has occurred within their lineage (red). Scale bar = 50um.
Link: https://www.youtube.com/watch?v=ZxywQ7LaihI&t=20s

**Video S2 | Single Cell Proliferation and Colony Expansion Heterogeneity.** A sequence of colourised binary masks with segmented individual cells (grey) on background (black), highlighting cell proliferation throughout the duration of a representative 1193 frame-long movie (~80 hours). Highlighted are the founder cells and progeny corresponding to slow (orange), medium (blue) and fast (green) dividers. Scale bar = 50um.
Link: https://www.youtube.com/watch?v=gScvX89JeYQ

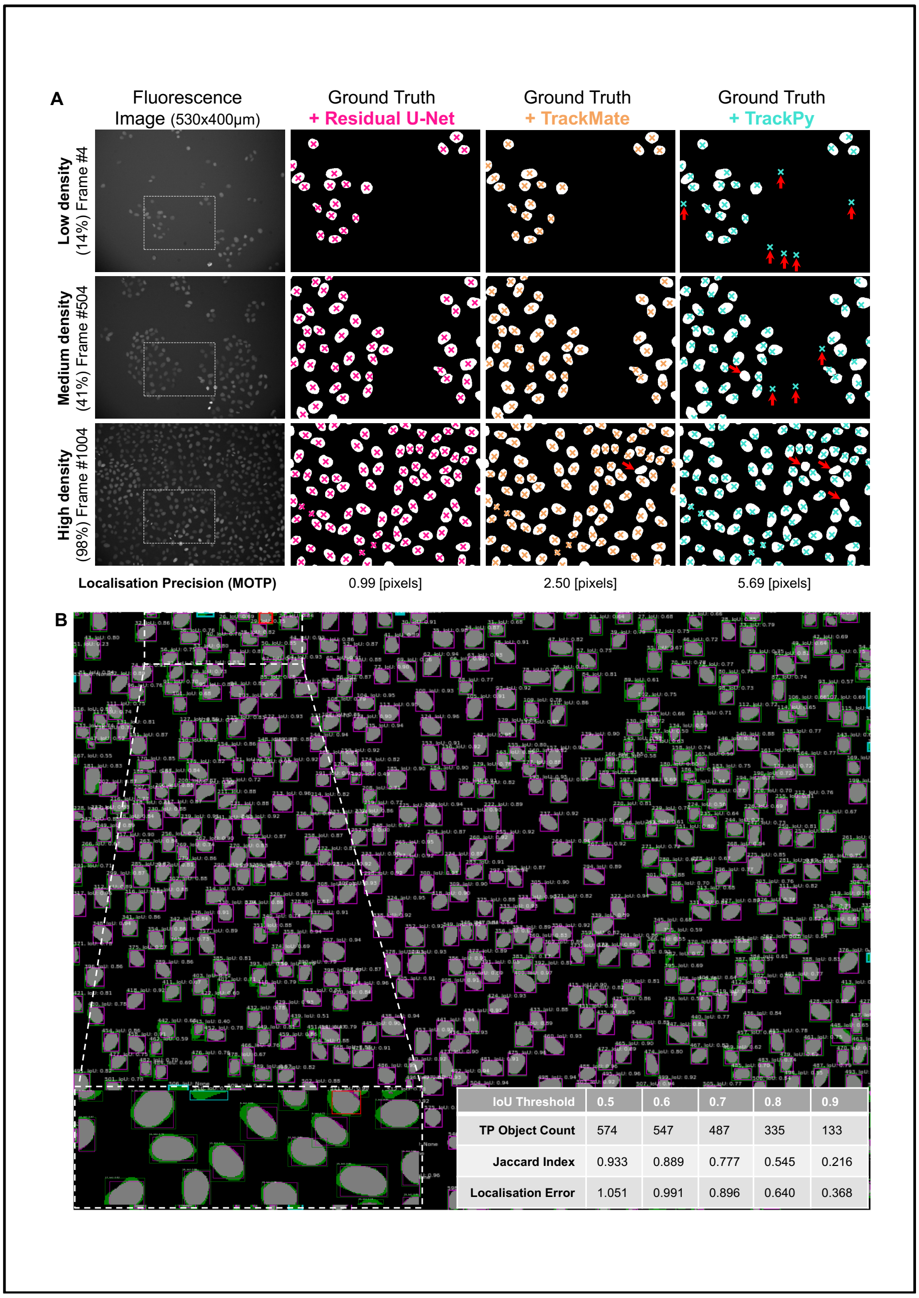


**Figure S1 | Cell Identification Precision by 3 Different Cell Detectors. A.** Comparison of Residual U-Net (pink), TrackMate (beige) and TrackPy (cyan) algorithms in detecting and localising cells in three representative fields of view with low, medium and high cell density. Overlaid are the cropped regions of ground truth segmentation masks with cell centroid markers with listed localisation scores below. Red arrows indicate the presence of false-negative (downward right) and false-positive (upward facing arrow) cell detection errors. **B.** Detailed overview of per-object bounding boxes and IoU scores map of high cell density field of view. Cell area and bounding box color-coding (green, ground truth only; magenta, U-Net prediction only; grey, match between true and predicted area) highlights cell detection accuracy, with different types of cell detection errors (cyan, false negative; red, false positive) depicted in zoomed-in region (lower left). Summary of Residual U-Net performance on the representative frame with strict IoU thresholding is described in accompanying table (lower right). Overall, we report the IoU of 0.802, Jaccard index of 0.975 and pixel identity correctness of 0.874 for all 3 tested fields of view on 869 objects with at least 1 pixel overlap between true and predicted area. TP, true positive; IoU, intersection over union.

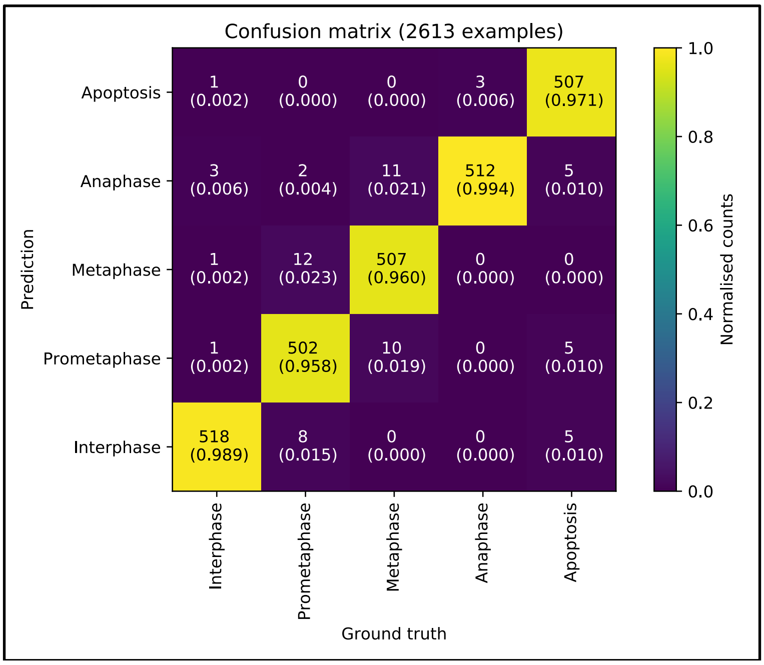


**Figure S2 | Confusion matrix for cell state labelling.** Confusion matrix for the testing dataset (n=2613 samples, ~500 examples per class) of brightfield and fluorescence cell image crops for the CNN model with normalised. Shown is the matching of human annotations vs. the CNN-generated annotations, showing the per-class label recall (rows) and precision (columns).

**A**

| Benchmarking Metric | bTrack | TrackMate | TrackPy |
|---|---|---|---|
| Localisation Accuracy (MOTA) | 97.66% | 97.54% | 73.77% |
| Mitotic Branching Correctness (MBC) | 0.87 (with penalty) | 0.42 (with penalty) | --- |
| | 0.87 (penalty-free) | 0.26 (penalty-free) | --- |
| Leaf Retrieval Score (LRS) | 0.78 (all leaf cells) | 0.14 (all leaf cells) | --- |
| | 0.84 (until last frame) | 0.29 (until last frame) | --- |
| Target Effectiveness – Tracker Recall (TE) | 90.55% (no stitching) | --- | --- |
| | 98.88% (subtrees) | --- | --- |
| | 99.83% (branches) | --- | --- |
| Track Purity – Tracker Precision (TP) | 99.72% (no stitching) | --- | --- |
| | 99.74% (subtrees) | --- | --- |
| | 99.75% (branches) | --- | --- |
| Stitching Actions | 8 subtree additions | --- | --- |
| | 1 subtree swap | --- | --- |
| | 25 branch additions | --- | --- |

**B**

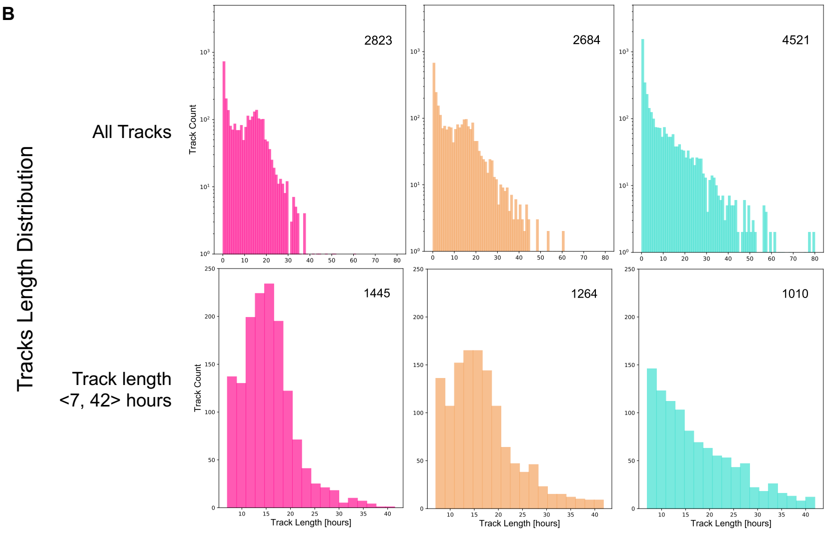


**Figure S3 | Benchmarking metrics amongst tracking pipelines. A.** Comparison of multiple benchmarking metrics to score the performance of *bTrack*, TrackMate and TrackPy algorithms with respect to single-cell trajectory following and lineage tree reconstruction. Due to poor performance of the TrackMate lineage tree reconstruction, target effectiveness and tracking purity scores were not calculated. TrackPy algorithm scoring is only listed for MOTA score as the algorithm is not designed to detect track splitting events and account for cell divisions. **B.** Histograms of track length distributions detected in all tracks (top) and tracks with typical cell cycle length (bottom) across the three tracking pipelines.

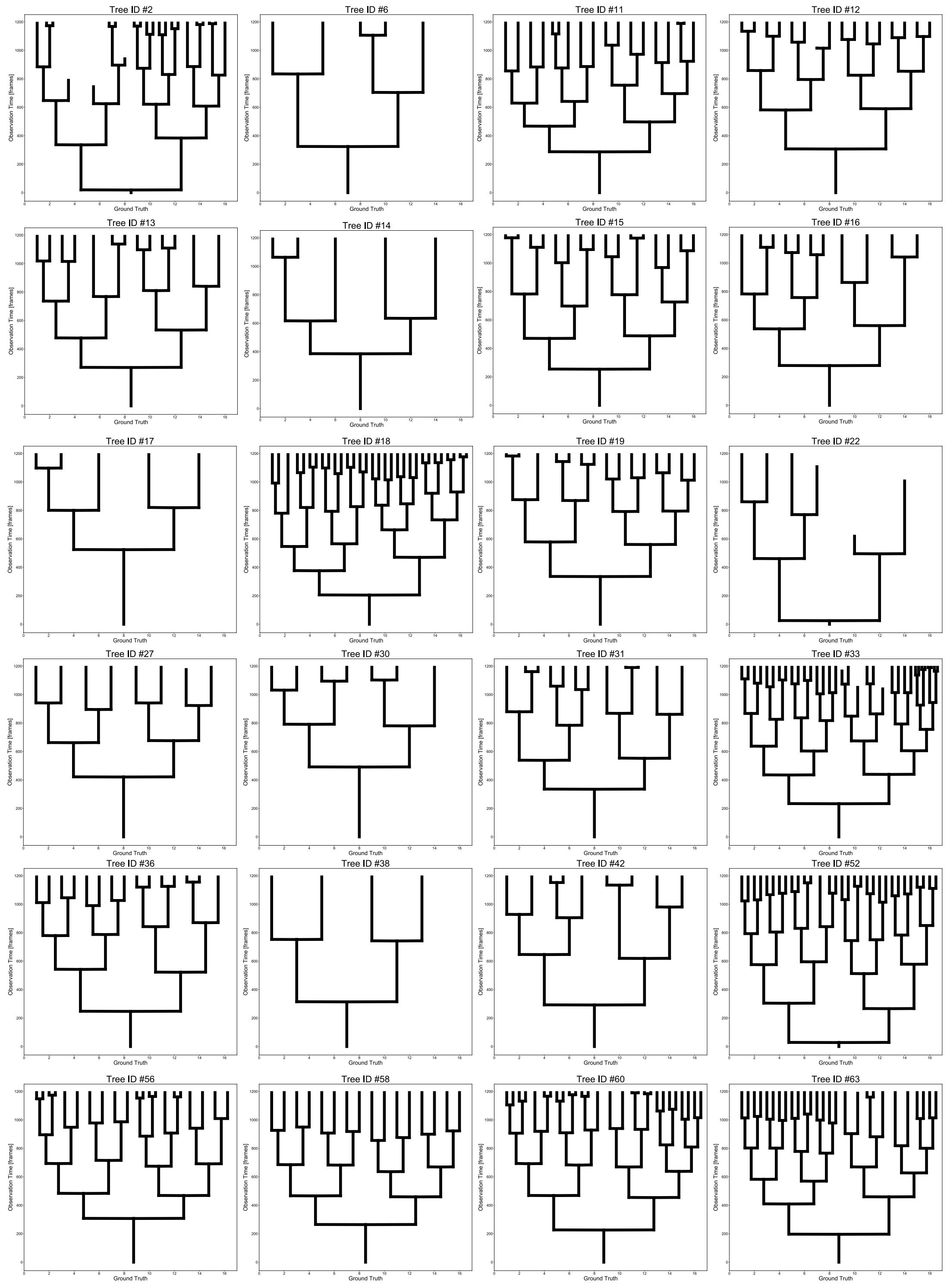


**Figure S4 | 2D Tree Representations of 24 Human Annotated Cell Lineages of a Representative Movie.** Y-axis represents the time elapsed from start of time-lapse imaging. Vertical lines correspond to cell cycle duration of the particular cell. Horizontal lines represent track splitting (cell division) events.

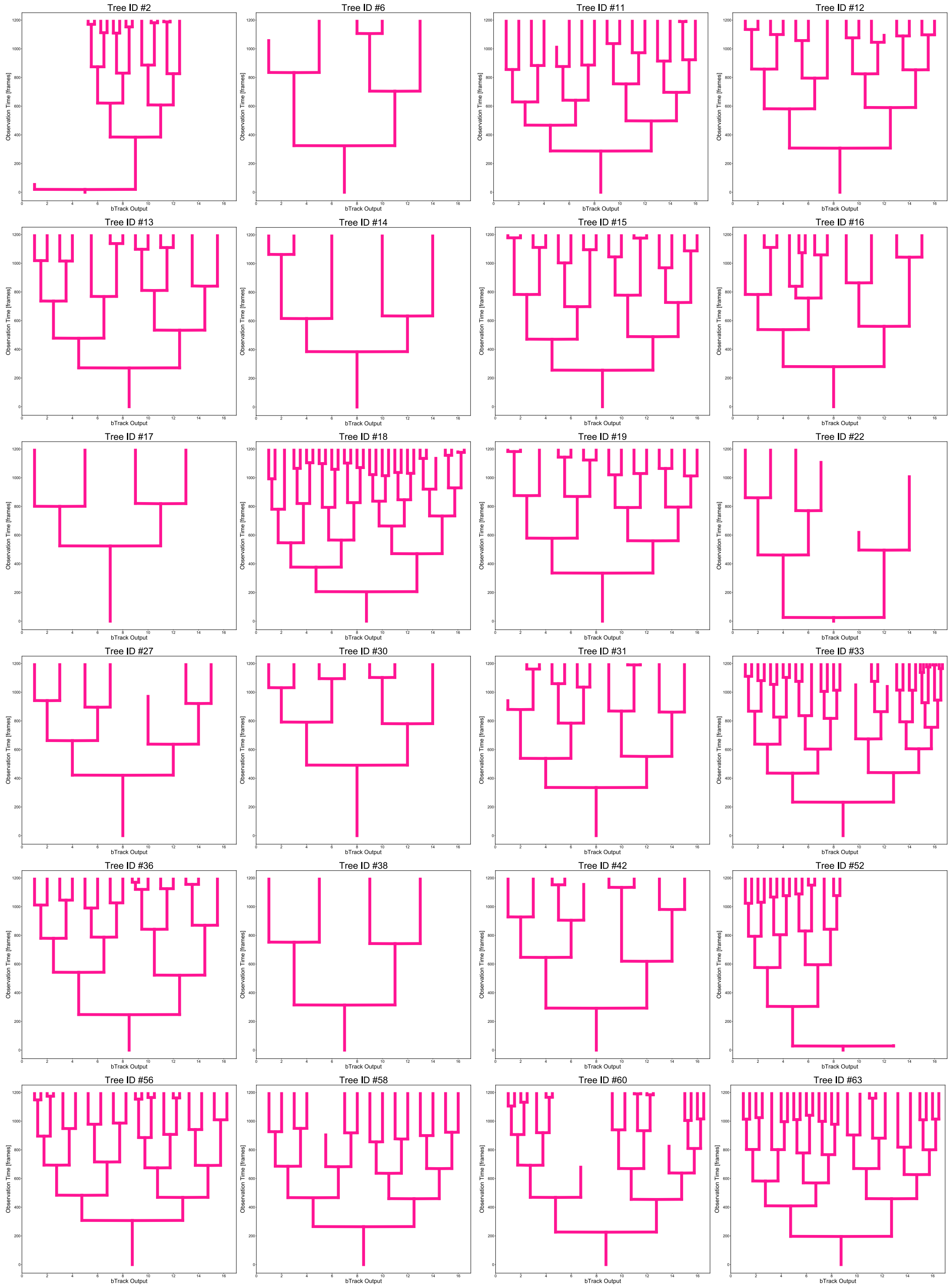


**Figure S5 | 2D Tree Representations of 24 Automatically Reconstructed Cell Lineages from a Representative Movie by our custom-designed *bTrack* pipeline.** Y-axis represents the time elapsed from start of time-lapse imaging. Vertical lines correspond to cell cycle duration of the particular cell. Horizontal lines represent track splitting (cell division) events.

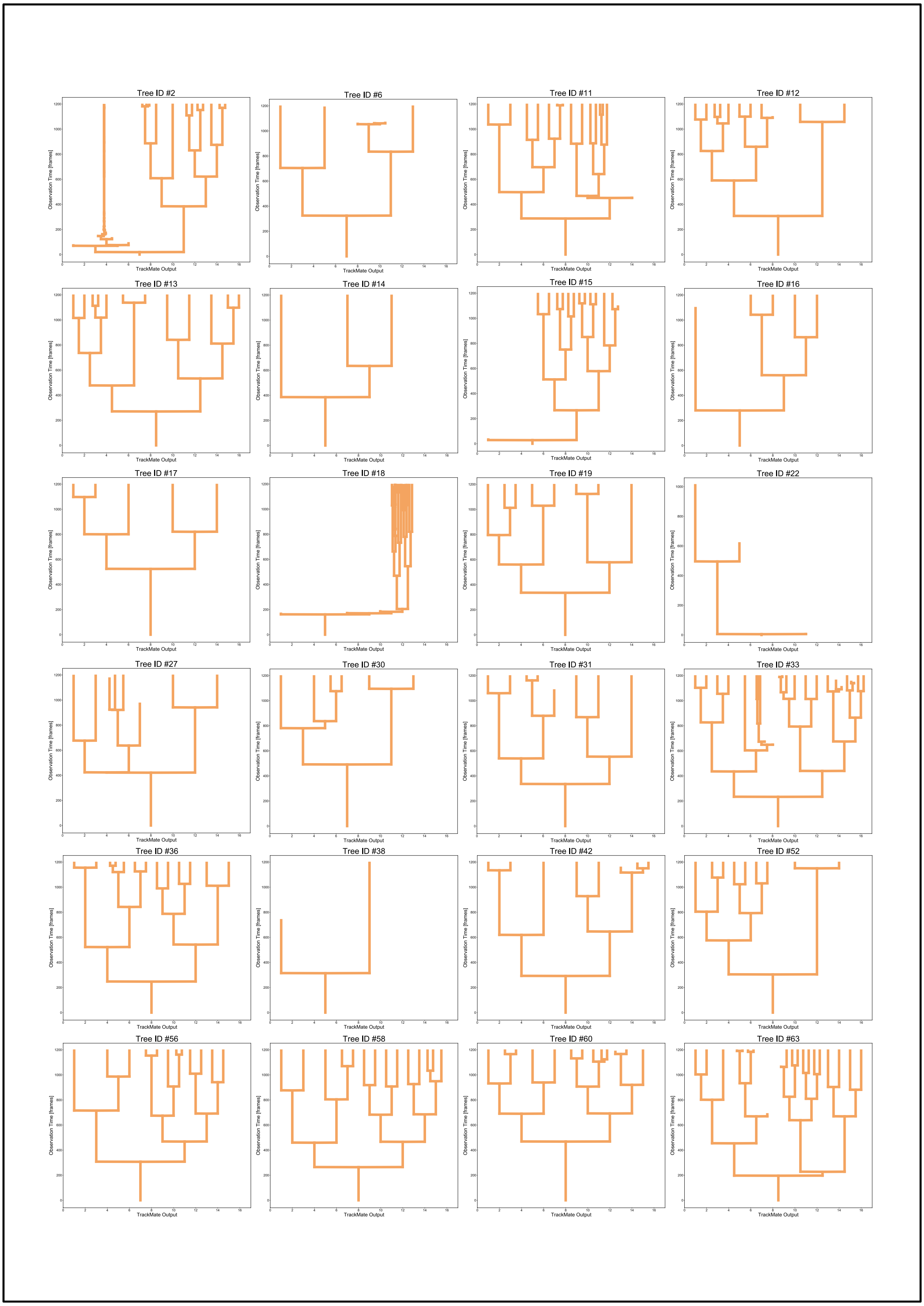


**Figure S6 | 2D Tree Representations of 24 Automatically Reconstructed Cell Lineages from a Representative Movie by a particle-tracking tool, *TrackMate*.** Y-axis represents the time elapsed from start of time-lapse imaging. Vertical lines correspond to cell cycle duration of the particular cell. Horizontal lines represent track splitting (cell division) events.

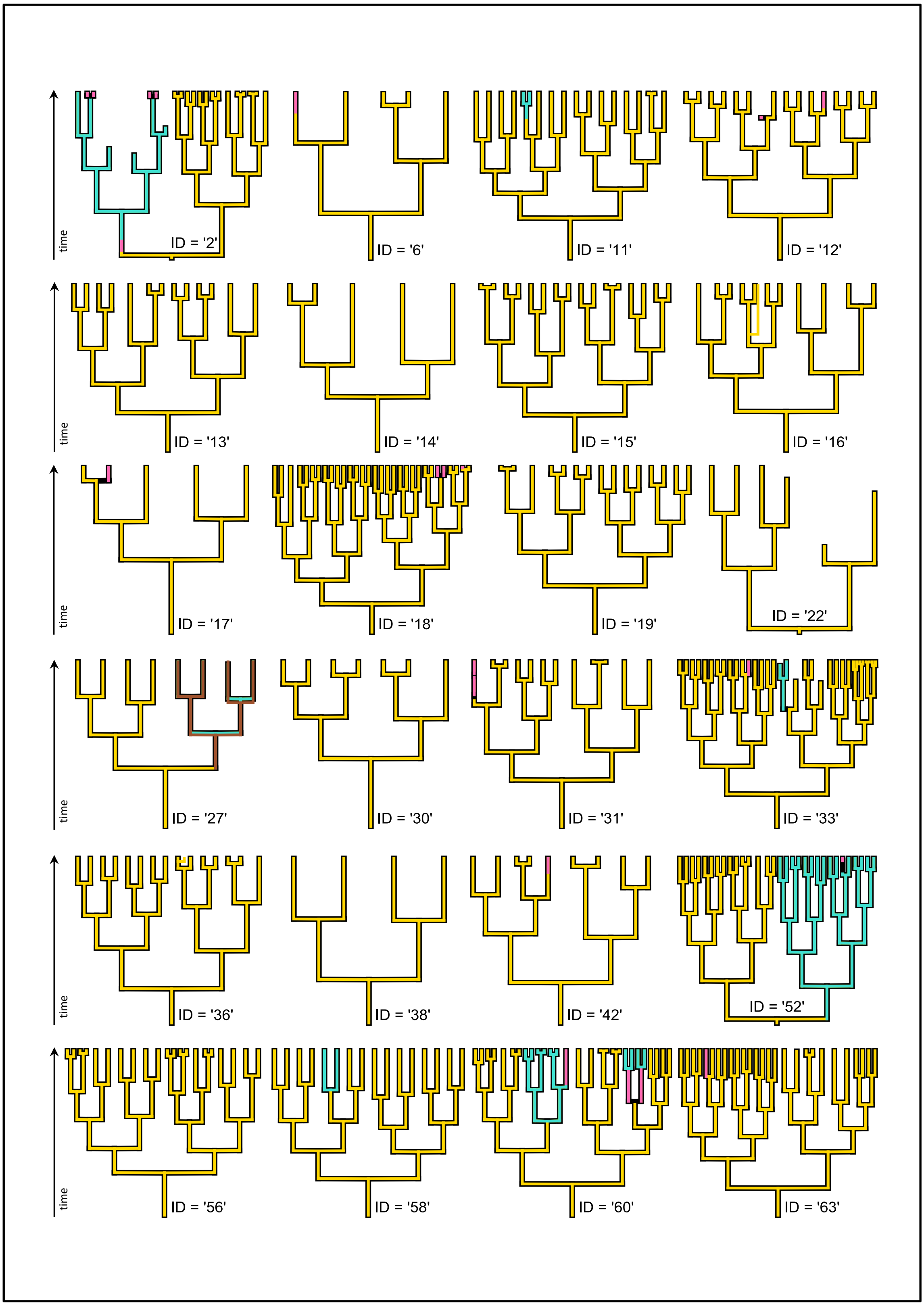


**Figure S7 | Visual Overview of Tree Re-Assembly.** Highlighted are the regions of ground truth trees (black thick background) which were correctly recapitulated by *bTrack* (gold branches). Upon branch breakage, two types of assembly actions were applied: subtree attachment (cyan), where the branch underwent further splitting, or branch attachment (pink), where the track did not further branch. In total, out testing tree pool comprised 8 perfectly tracked trees (ID: 13, 14, 15, 19, 22, 30, 38, 56), with 7 trees requiring one or more subtree stitch (ID: 2, 11, 27, 33, 52, 58 and 60). In case of tree #27, the right subtree was falsely associated with the tree (brown) and was swapped accordingly.

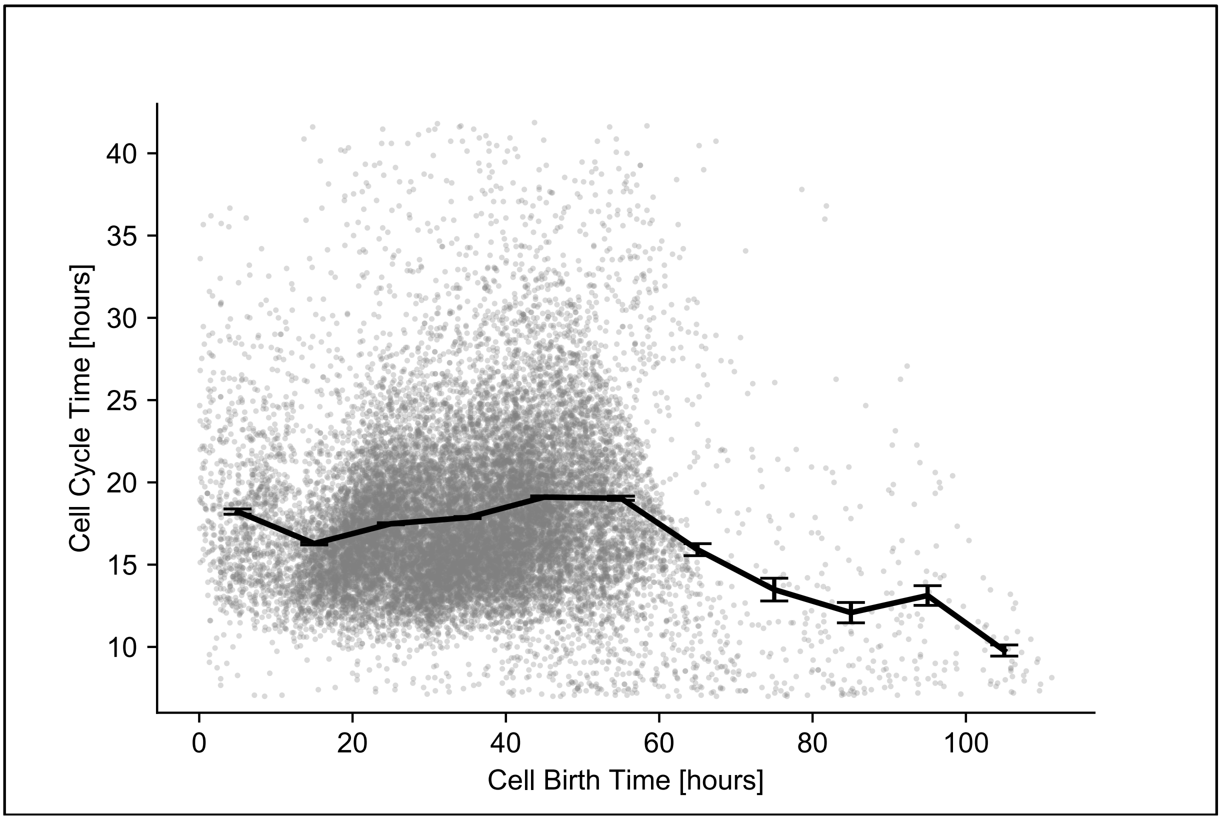


**Figure S8 | Cell cycle heterogeneity does not occur due to the time of cell birth.** Cycle lengths of fully resolved cells show no trend over the duration of live-cell imaging with respect to cell birth time relative to start time. For the first 60 hours of time-lapse imaging, the mean intermitotic time was determined to be constant. Due to variable durations of movie time-lapses, ranging between 64 and 120 hours, the mean cell cycling duration appears to decrease at longer imaging durations as only cells born with progressively shorter cycling lengths are sampled. Error bars indicate standard error of the mean calculated for 10-hour bins.
